# Physiological Markers of Auditory Situational Awareness in Complex Spatialised Scenes

**DOI:** 10.64898/2026.05.27.728177

**Authors:** Jakub Sztandera, Katarina C. Poole, Martha Shiell, Lorenzo Picinali, Maria Chait

## Abstract

Detecting changes in acoustic environments is essential for situational awareness. It remains unclear whether the spatial location of such changes modulates automatic orienting and arousal mechanisms. We measured pupil dilation, pupil dilation rate, and microsaccade rate while listeners (n=25) heard complex, spatialized auditory scenes rendered over headphones using individualized HRTFs. Participants were naïve to the critical manipulation: the appearance of a new source from one of five locations: front, left, right, back, or above. A subsequent localization task assessed perceptual spatial uncertainty.

Behaviorally-irrelevant source appearances elicited a cascade of ocular responses. Microsaccadic inhibition emerged from ∼85ms after change onset, and was broadly comparable across locations, suggesting a location-invariant early orienting response to auditory change. Pupil dilation rate increased from ∼200ms, followed by a phasic pupil dilation response from ∼400ms, indicating engagement of arousal-related systems. Pupil responses were modulated by source location: changes from front/left/right elicited larger dilation than changes from above, with back responses showing a similar but weaker reduction. Behavioral localization revealed substantial confusion for front/back/above locations. However, this did not mirror the physiological data, as front sources elicited pupil responses comparable to lateral sources.

These findings demonstrate that complex auditory scene changes recruit oculomotor and autonomic systems even outside the focus of task relevance. They further suggest a dissociation between early, location-invariant attentional capture indexed by microsaccadic inhibition and later, location-sensitive arousal indexed by pupil dilation. Spatial biases in auditory situational awareness therefore appear to emerge after initial change detection, shaping arousal and behavioral performance rather than the earliest orienting response.

## 1. Introduction

Detecting changes in the acoustic environment is critical for situational awareness and adaptive behaviour. In complex natural scenes, listeners must rapidly identify deviations from ongoing sound patterns, such as those accompanying the appearance or disappearance of sound sources, in order to reorient attention toward potentially relevant events. A substantial body of research shows that the auditory system is highly sensitive to such deviations (Aman et al., 2021; Constantino et al., 2012; de Kerangal et al., 2021; Demany et al., 2017; Gregg & Samuel, 2008; Irsik et al., 2016; Pavani & Turrato, 2008), with neural and physiological signatures emerging within a few hundred milliseconds of change onset, even when listeners are not actively engaged in a change detection task (Huviyetli & Chait, 2026; Sohoglu & Chait, 2016a,b). These responses are widely interpreted as reflecting automatic change-detection mechanisms that operate pre-attentively as part of a “non-target” listening mode, enabling the system to monitor the acoustic scene even when attention is directed elsewhere.

Several physiological measures provide complementary insights into the dynamics of auditory change detection. Early cortical M/EEG responses to scene changes typically emerge approximately 50–200 ms after change onset (Poole et al., 2025a; Sohoglu & Chait, 2016a,b) and are thought to reflect automatic change-detection processes in the auditory cortex.

Sohoglu and Chait (2016a,b) first characterized these responses using simple artificial acoustic “scenes” composed of multiple streams of pure tones, in which streams abruptly appeared or disappeared partway through the scene. More recently, Poole et al. (2025a) extended this paradigm to more complex, spatialized, scenes consisting of multiple wideband sound sources.

These stimuli were designed to approximate the temporal fluctuation dynamics of natural acoustic environments while removing semantic attributes associated with identifiable sound sources. In both studies, EEG responses to scene changes were observed in naïve listeners performing an unrelated decoy task. In the simpler tone-based scenes used by Sohoglu and Chait (2016a,b), responses emerged from approximately 50 ms after the change, whereas in the more complex scenes used by Poole et al. (2025a), responses appeared later, from around 200 ms post-change.

Additional indexes of processing are available through measurement of two different ocular responses, pupil size and microsaccades, which reflect the operation of circuitry related to deployment of arousal and attentional capture (Huviyetli & Chait, 2026). Phasic pupil dilation responses are associated with arousal and orienting to salient events and are thought to reflect activity in neuromodulatory systems involved in attention and alerting (Sara & Bouret, 2012).

Pupil size is widely used as a proxy for locus coeruleus (LC) activity: baseline PD reflects tonic LC activity associated with general alertness, whereas transient dilations index phasic LC activations that signal rapid arousal to novel stimuli (Aston-Jones and Cohen, 2005; Joshi et al., 2016; Wang and Munoz, 2021). Microsaccades are small, involuntary fixational eye movements that are thought to support the automatic exploration of the visual environment (Martinez-Conde et al., 2006; Otero-Millan et al., 2008). The presentation of novel stimuli, across sensory modalities and not limited to vision, elicits a brief suppression of microsaccades (Engbert & Kliegl, 2003; Rolfs et al., 2008). This microsaccadic inhibition (MSI) is thought to reflect attentional reorienting and the allocation of processing resources to a new event. Indeed, the magnitude and time course of MSI are shaped by stimulus salience and the observer’s attentional state (Bonneh et al., 2014; Huviyetli & Chait, 2026; Kadosh & Bonneh, 2022; Liu & Chait, 2026; Roberts et al., 2019; Rolfs et al., 2008; White & Rolfs, 2016; Zhao et al., 2024).

Taken together, these findings suggest that MSI reflects a rapid and adaptive reorienting mechanism that transiently interrupts ongoing (visual) processing in order to prioritize unexpected inputs and support appropriate behavioral responses. Quantifying MSI to new events therefore provides an indirect measure of whether, and to what extent, those events triggered reorienting processes.

Recently, Huviyetli and Chait (2026) demonstrated a robust ocular responses to behaviourally irrelevant scene changes. In their study, artificial acoustic scenes composed of pure-tone streams were presented with occasional changes introduced by the appearance of a new source. Participants remained naive to these changes and performed an unrelated decoy task throughout. An MSI response emerged approximately 70 ms after the change and reached peak inhibition at around 250 ms. It was followed by a phasic pupil dilation (PD) beginning roughly 400 ms post-change. These responses were interpreted as reflecting bottom-up attentional capture and arousal, respectively, elicited by unexpected changes in the acoustic scene despite their lack of behavioral relevance.

The conceptual association between arousal and attention is supported by overlapping neural circuitry. The control of PD and MS depends on interconnected networks, including reciprocal connections among the LC, frontal eye fields (FEF), and superior colliculus (SC), regions implicated in the generation and control of microsaccades (Hafed et al., 2009; Joshi & Gold, 2020; Wang et al., 2020). Nevertheless, growing evidence indicates that PD and MS reflect at least partially dissociable processes (Contadini-Wright et al., 2023; Huviyetli & Chait, 2026; Liu & Chait, 2026; Zhao et al., 2024). For example, MS effects tend to be restricted to temporal windows in which attention is actively deployed, such as the reduction in MS rate during presentation of behaviourally relevant keywords (Contadini-Wright et al., 2023; Liu & Chait, 2026). By contrast, PD responses are often more sustained, consistent with longer-lasting arousal-related activity. In line with this distinction, PD and MS frequently exhibit different patterns of modulation. Huviyetli and Chait (2026) reported a facilitative effect of predictability on MS but not PD, whereas Liu and Chait (2026) found an effect of active ignoring on PD but not MS. They interpreted this dissociation as evidence that MS rate is specifically shaped by attentional allocation, whereas PD reflects a more general arousal mechanism. Taken together, these findings suggest that PD and MS provide complementary indices of the engagement of arousal and attentional systems during task performance.

In the present study, building on the paradigm developed by Poole et al. (2025a), we examined PD and MS responses to scene changes in complex, spatialized acoustic scenes. Spatial information is a core component of auditory scene analysis (van der Heijden et al., 2019) and is central to situational awareness. In principle, the location of an unexpected sound may influence the extent to which it captures attention and guides subsequent orienting behaviour (Bayram et al., 2025; Krishnamurthy et al., 2017; Spence, Lee, & Van der Stoep, 2017). However, relatively little is known about how the spatial location of events modulates their capacity to attract attention. Furthermore, human’s ability to process spatial cues and determine the source’s location is often not reliable. A well-known limitation is front–back confusion, whereby sources in front of and behind the listener are mislocalized due to ambiguity in interaural cues along the cone of confusion, particularly when spectral and dynamic cues are weak (Carlile & Pralong, 1992). This raises a key question for the present work - whether spatial ambiguity constrains the processing of scene changes. Specifically, it remains unclear whether such uncertainty affects early, automatic orienting responses, or emerges only at later stages of perceptual inference and response selection.

In their EEG study, Poole et al. (2025a) examined whether the location of a new source appearing partway through an acoustic scene modulated neural responses to change. In a sample of 60 participants, they found no effect of change location: Early EEG responses to changes originating from the front, left, right, above, or behind were comparable. However, behavioural experiments using the same stimuli, as well as follow-up work by Poole et al. (2026), revealed slower responses to changes occurring behind and above the listener. This pattern was interpreted as reflecting the emergence of a perceptual bias toward the frontal hemifield. Commensurate spatial biases have been reported elsewhere. For example, attentional biases toward the direction of gaze have been observed in both auditory and multisensory contexts (Best, Boyd, & Sen, 2023; Pomper & Chait, 2017), consistent with the idea that the perceptual system upweights information arising from behaviourally relevant regions of space, particularly those aligned with current gaze. At the same time, biases favouring the rear space have also been reported. Bodnár, Mock, and Golob (2025), for instance, found faster and more accurate responses to sources originating behind blindfolded listeners. Similarly, Baek et al. (2025) showed that blindfolded individuals tend to mislocalize two spoons being struck directly in front of them as originating from behind. These findings have been interpreted as reflecting a survival-related bias to prioritize potential threats arising from outside the field of view.

A key step toward understanding how these spatial biases arise is to determine whether they emerge during automatic processing, when listeners are not actively attending to spatial information. Previous work examined this question using EEG. In the present study, we extend this approach by examining how automatic auditory change detection interfaces with neural systems governing arousal and attention. Specifically, we focus on autonomic ocular responses to determine whether the location of a newly appearing source within a complex, spatialized auditory scene engages physiological systems associated with MS-linked attentional capture and PD-linked arousal, even when the change is behaviourally irrelevant. We test whether the spatial biases previously observed in behavioural localization (Poole et al., 2025a; 2026) are also expressed in ocular markers of change detection, reflecting the downstream recruitment of attention and arousal networks which are activated following the change detection process to support rapid orienting and adaptive responses (e.g., fight or flight).

If sensitivity to spatial reliability emerges early in the processing stream, it should be evident in early physiological responses, potentially delaying or reducing responses from certain spatial locations. In particular, modulation of MSI, emerging from approximately 80 ms after event onset, provides an index of attentional capture and stimulus salience. Arousal-related processing can be assessed through pupil dilation rate and pupil dilation amplitude, with effects expected from approximately 400 ms onward. By contrast, if these spatial biases emerge only at later stages of perceptual evaluation or response selection, then automatic ocular responses should not vary systematically with source location.

## 2. Methods

The study consisted of three parts: (a) a head-related transfer (HRTF) measurement, (b) an eye tracking experiment whilst participants listen to spatialized scenes over headphones and perform an decoy gap detection task, and (c) a behavioral localization task over loudspeakers and headphones.

### 2.1. Participants

All participants reported normal hearing, normal or corrected-to-normal vision (sphere prescription no greater than +/-3.5 D), and no history of neurological disorders. The experiment was approved by the UCL Research Ethics Committee (Project ID: 1490/011). Individual HRTFs were obtained as part of the SONICOM project (approved by the Imperial College Ethics Committee; SETREC number: 6533981). Participants provided informed consent and were compensated for their time.

We aimed to achieve N>= 25 based on previous pupillometry research on auditory change detection(Huviyetli & Chait, 2026; Zhao et al., 2024). 36 participants were recruited in total. Data from 3 participants were excluded due to technical issues with audio presentation; data from 7 participants were excluded due to excessive missing pupillometry samples caused by blinks, fatigue, or tracking loss (see Data Preprocessing and Statistical Analysis). One further participant was removed due to poor performance on the decoy task. The final sample comprised 25 participants (19 female; mean age = 24.12 years). This data was used in the analysis of the eye tracking experiment and HRTF-based behavioral localization task. Partway during data collection we extended the localization task to include a free-field condition which was completed by 18 participants (11 female; mean age = 25.33 years).

### 2.2. Stimuli

The stimuli used in the eye tracking experiment and localization test were identical to those in Poole et al. (2025a). These were ‘chimeric’ sounds created by extracting the temporal envelope from naturalistic modulator sounds (e.g., environmental recordings) and applying it to synthesized carrier sounds to create each individual source (for detailed description see Poole et al., 2025a). Using these stimuli allowed us to minimize any confounding effects that might have been caused by semantic content, whilst ensuring they possess naturalistic-like properties and broadband spatial cues (see Figure 1A).

**Figure 1.**
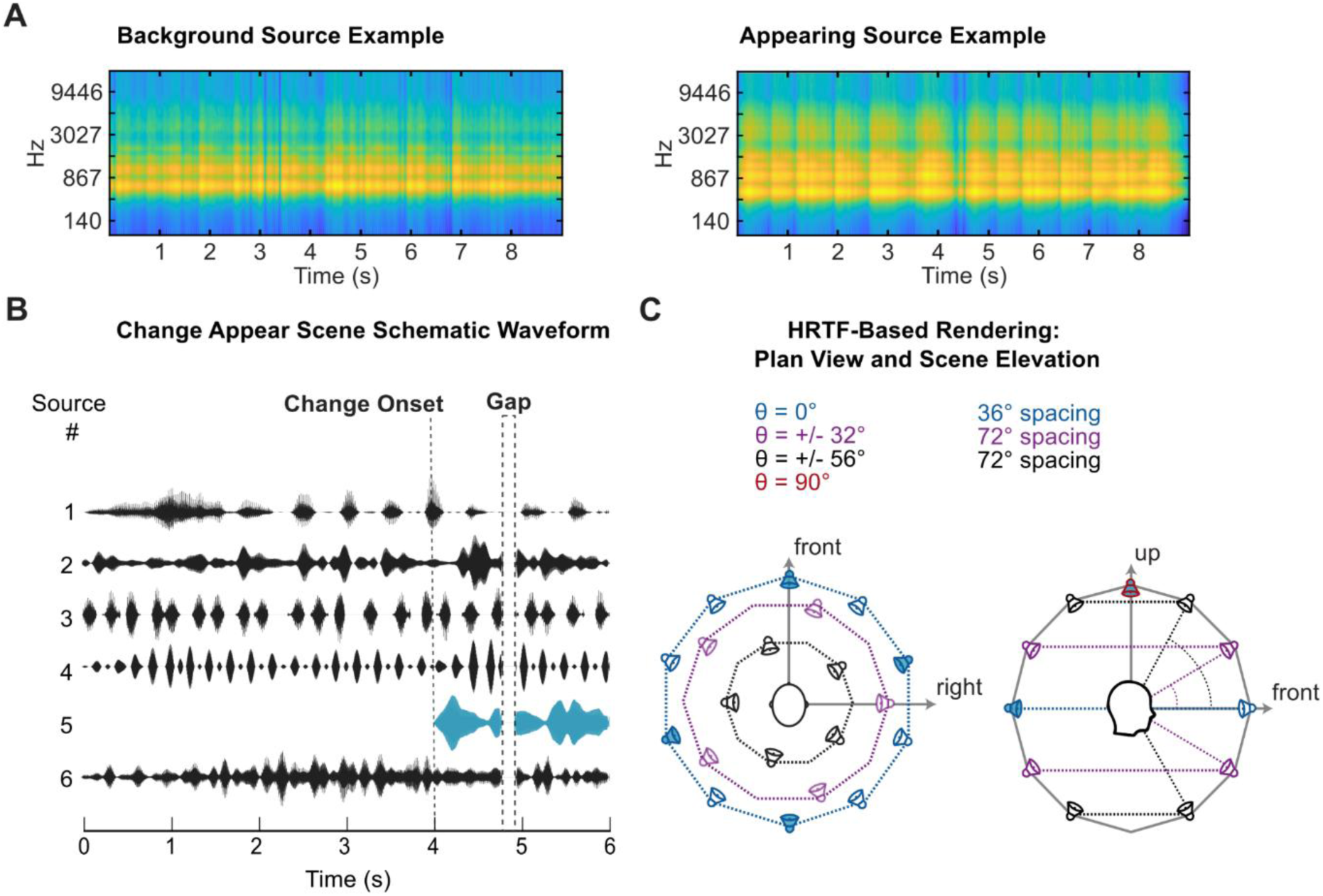
Stimuli and apparatus. **[A]** Representative ERB-weighted spectrograms of background and appearing stimuli. **[B]** Schematic of an example trial showing a multi-source Change Appear scene illustrating the onset of an appearing source (teal) at 4 s and the task-related silent gap. The background source waveforms (1-4, 6) are presented in black. **[C]** Schematic of the possible source locations in the HRTF-rendered scenes in plan view (left) and side elevation (right), coloured by elevation angle (teal = 0°, purple = ± 32°, black= ± 56°). Speakers filled in blue show possible appearing source locations and background sources could be presented from any available source location.

As in Poole et al. (2025; 2026) a total of 15 sources were divided into five possible appearing sources and 10 background sources. The background sources were then combined into 6 s multi-source “scenes” that contained five background sources. Unlike Poole et al (2025; 2026), in the eye tracking experiment the stimuli were rendered binaurally, i.e., using individualised HRTF filters to produce virtual spatialised sounds over headphones rather than presented over a free-field loudspeaker array. To enable individualised HRTF-based rendering, participants first attended an HRTF measurement session conducted in an acoustically treated room at Imperial College London, at least a few days prior to the eye tracking and localisation experiments (for full details of materials and procedure see Engel et al. 2023; Poole et al. 2025b). Each source was convolved offline with the participants’ individual HRTF (SOFA format) using the *py3dti* binaural renderer (Pauwels, 2023) to simulate a three-dimensional auditory source at a given location. This was at a simulated distance of 1.5 m consistent with the HRTF measurement distance. The resulting binaural signals for all sources within a scene were then summed and combined into a single stereo file at a sample rate of 48 kHz, such that when listened to over headphones each source was perceived at its virtual spatial position.

The virtual locations of the sources were identical to real speaker locations in the 31-speaker array in Poole et al. (2025a; illustrated above in Figure 1C). Appearing sources were assigned to five possible locations: four were at ear level (elevation = 0) and one was directly above the listener (elevation = 90). The four ear level locations were Front, Left, Back, and Right, corresponding to counterclockwise azimuths of 0°; 72°; 180°; 252°, respectively (see Figure 1C). For the placement of background sources, we created a balanced distribution by dividing the 31 positions into six equal-sized radial segments. Background sources were pseudo-randomly allocated to five unoccupied segments.

During the eye tracking experiment, participants listened to these stimuli, presented with an interstimulus interval of 5 s, whilst performing a decoy gap detection task. Approximately 10% of the stimuli contained a silent gap (150 ms; onset and offset shaped with a 20 ms raised-cosine ramp) inserted randomly between 1 to 5 s post-onset. The conditions of interest were the appearance of a new source (“Change appear”; “CA”) vs no change (“NC”) and location of the appearing source (Front, Left, Right, Back, Above). Listeners were presented with 30 trials per condition. The experiment was divided into ∼10 minutes-long blocks each containing 5 trials per condition (for a total of 50 trials) and 4–6 gap containing decoy-task trials (pseudorandomized to reduce expectancy effects).

### 2.3. Procedure

The eye tracking and localization task were conducted at the UCL Ear Institute in a dimly lit triple-walled sound–attenuating booth (IAC). During eye tracking, participants were seated in front of a 24-inch BENQ XL2420T screen (resolution: 1920x1080, refresh rate: 60Hz), with their head placed on a chinrest. They were instructed to fix their gaze on a 50x50px black fixation cross at the center of a light grey screen with a luminance of 63.34 cd/m². Eye position and pupil size data were collected at a rate of 1000 Hz using an infrared eye-tracking camera (EyeLink 1000 Desktop Mount, SR Research) that was located at a distance of 62 cm. Participants listened to scenes rendered using HRTFs through Sennheiser HD558 headphones and a Roland Tricapture 24-bit 96 kHz soundcard. A standard five-point calibration was performed at the beginning of each block. Before each trial, gaze position was checked. The trial would not commence unless the gaze was within 50 px of the fixation cross. This detection was automated, and fed back to the participants by turning the fixation cross red. Participants were naive to the experimental manipulation. They were instructed to listen to the presented acoustic scenes and report (with a key press) the presence of gaps in the audio. This task maintained attentional engagement and served as a decoy from the main experimental manipulation (appearance events). To maintain engagement, feedback was provided after each trial and at the end of each block.

Lastly, participants completed the behavioral localization task. The test consisted of 25 CA scenes (five scenes per location condition). Participants were instructed to listen to each scene and indicate, on a visual display, the perceived location of the change and their confidence in their judgment on a five-point Likert-type scale, from 0 (complete guess) to 4 (very confident). Participants completed the localization task in either one or two rendering conditions. Ten participants completed the HRTF condition only, presented over headphones using the same individualised HRTF-based rendering as the eye tracking experiment described above. While 18 participants completed both the HRTF and free-field conditions in counterbalanced order.

In the free-field condition, scenes were delivered through a custom (2.2 x 2.2m) array of eight loudspeakers (Genelec, 8010A) with participants seated at the center of the loudspeaker array and facing an LG Flatron 24EN43V-B display (24.0″ diagonal, 1080p). Four loudspeakers were located at ear level, with azimuths of 0°, 90°, 180°, and 270° with the corresponding approximate distances from the listener position of 1.4, 1.36, 1.4 and 1.4 m, respectively. Four further loudspeakers were located at approximately 37° elevation relative to participants’ ears, at the 45°, 135°, 225°, and 315° azimuths. The corresponding approximate distances from the listener position were 1,13, 1.35, 1.18, and 1.22 meters, respectively. To approximate the source positions of the 31-loudspeaker array as in our previous work (Poole et al., 2025a), phantom sources were created (Theile & Plenge, 1976), in Max/MSP (v8.0, Cycling ’74) using the IRCAM Spat package (Carpentier et al., 2015), which performed amplitude panning across loudspeakers neighbouring the desired virtual position (Pulkki, 1997). Auditory presentation was controlled via a MOTU 24Ao audio interface. The assignment of background sources in the localization test was handled identically to the main experiment. Appearing sources were presented as real loudspeaker sources for ear-level locations and as phantom sources for the Above condition. Background source assignment was handled identically to the main experiment.

### 2.4. Data Preprocessing and Statistical Analysis

#### 2.4.1. Behavioral analysis

Gap detection task: Key presses that occurred <2 s following a target gap, were designated as hits; responses when no gap was present were designated as false alarms. Overall, participants made few false alarms (see Results); therefore, only hit rates were analysed. To ensure appropriate task engagement, data from participants who exhibited low performance (1 participant; see above) were excluded from further analysis. Trials containing a gap, or those in which participants produced a false alarm were removed from all ocular data analyses.

Localization test: Localization performance in the HRTF-rendered audio condition was assessed for all participants whose ocular data met inclusion criteria. Normality of localization accuracy and confidence score data was evaluated using Shapiro–Wilk tests. As these data violated normality assumptions, the global effect of source location on localization accuracy and confidence was analyzed using Friedman tests, followed by Bonferroni-corrected pairwise Wilcoxon signed-rank tests. For the subset of participants who completed both the HRTF and Free-Field localization conditions, localization accuracy was further analyzed to assess the effects of rendering method and source location. The effects were tested using a two-way repeated-measures ANOVA.

#### 2.4.2. Microsaccade rate

The microsaccade (MS) detection algorithm based on Engbert and Kliegl (2003) was applied to the horizontal continuous eye-movement data after removal of segments with partial or full eye closure. MS events were identified using the following criteria: (1) movement exceeding velocity threshold of λ = 6 times the median-based standard deviation of velocity distribution for each block, (2) one which exceeded that threshold for 5-100 ms, (3) with onset disparity across both eye data streams of <10 ms, (4) and separated by at least 50 ms from successive MS onsets. At the preprocessing stage the epochs contained the entirety of the scene presentation (-4 s : 2 s relative to change onset), and those with >50% of missing data were removed from the analysis. The resulting time series represented MS events as unit impulses (Dirac delta). MS rate was obtained by summing event time series across trials and normalizing by the number of trials and sampling rate. Next, we applied a causal smoothing kernel with a decay parameter of α = 1/50 ms, in line with the standard technique for computing neural firing rates (Dayan and Abbott, 2001; Rolfs et al., 2008). The time axis was shifted by the peak of the kernel to account for the kernel-induced temporal delay.

#### 2.4.3. Pupil dilation

Only left-eye data were analyzed. Samples in which the gaze deviated by more than 100 pixels from the fixation cross or contained partial or full eye closure (e.g., blinks) were marked as missing. To account for blink-edge artifacts, an additional 250 ms before and after each eye-closure event was also treated as missing. Missing segments were interpolated using a shape-preserving piecewise cubic interpolation method, and the pupil time series was then smoothed with a 150 ms Hanning window.

The analysis focused on change-evoked pupil responses. Trials were epoched from −500 ms to 2 s relative to change onset. A 50% exclusion criterion was applied: any epoch with more than half of its samples missing, or any participant with more than half of their trials missing in a given CA × Location condition, was excluded from further analysis. Additionally, outlier trials in which at least 20% of samples deviated by more than two standard deviations from the condition mean were removed. On average, four trials per condition were excluded per participant. The remaining trials were z-score normalized using the mean and standard deviation computed over the entire block, time-domain averaged, and baseline corrected using a 200 ms pre-change interval.

#### 2.4.4. Pupil dilation rate

We also analyzed the incidence of pupil dilation events (i.e. the points in time where the pupil begins to dilate), quantified as pupil dilation rate (PDR). Previous work has demonstrated that PDR correlates with spiking activity in three midbrain structures related to orienting and arousal: locus coeruleus and superior and inferior colliculi (Joshi et al., 2016). In line with previously used methods (e.g., Joshi et al., 2016; Zhao et al., 2024), pupil dilation events were identified as local minima found in smoothed continuous pupil data (150 ms Hanning window) that preceded at least 100 ms of continuous dilation. These events were modelled as Dirac functions (following the methods used to assess microsaccade rate) and analysed as described for the microsaccade data (above).

#### 2.4.5. Group-level analysis

Our aim was to evaluate the overall change detection response, measured via PD, PDR, and MS metrics, and how it varied by location. We used a nonparametric bootstrap analysis (Efron and Tibshirani, 1994). For condition comparisons, we generated repeated-measures difference time series by subtracting condition means; The difference time series were then randomly sampled (with replacement) to generate 1000 bootstrap samples. To classify segments containing significant differences between conditions, we identified time points where >95% bootstrap iterations fell consistently above or below 0. For PD, the analysis was conducted over the change-related period (from -500 ms before change onset to scene offset). In the analysis of MSI and PDR effects we focused on a predefined region of interest covering the first 500ms post change onset, which captures the typical latency of these responses. For the between-location comparisons, we controlled the likelihood of spurious effects by defining a significance threshold based on the longest contiguous significant baseline segment observed across analyses of difference waveforms generated from 100 iterations of NC trial sampling. We applied permutation-based bootstrap control analysis for FDR correction in PD response analysis by creating noise distributions through resampling NC data and comparing with the observed effect.

## 3. Results

### 3.1. Decoy gap detection task confirms engagement with auditory stimuli

First, we examined whether participants engaged correctly with the decoy gap detection task. As shown in figures 2A,B, most participants performed at or near ceiling on the decoy gap-detection task, with a mean reaction time of 791.8 ms. On average, participants produced fewer than three false alarms over the course of the session (max = 10; mode = 1).

**Figure 2.**
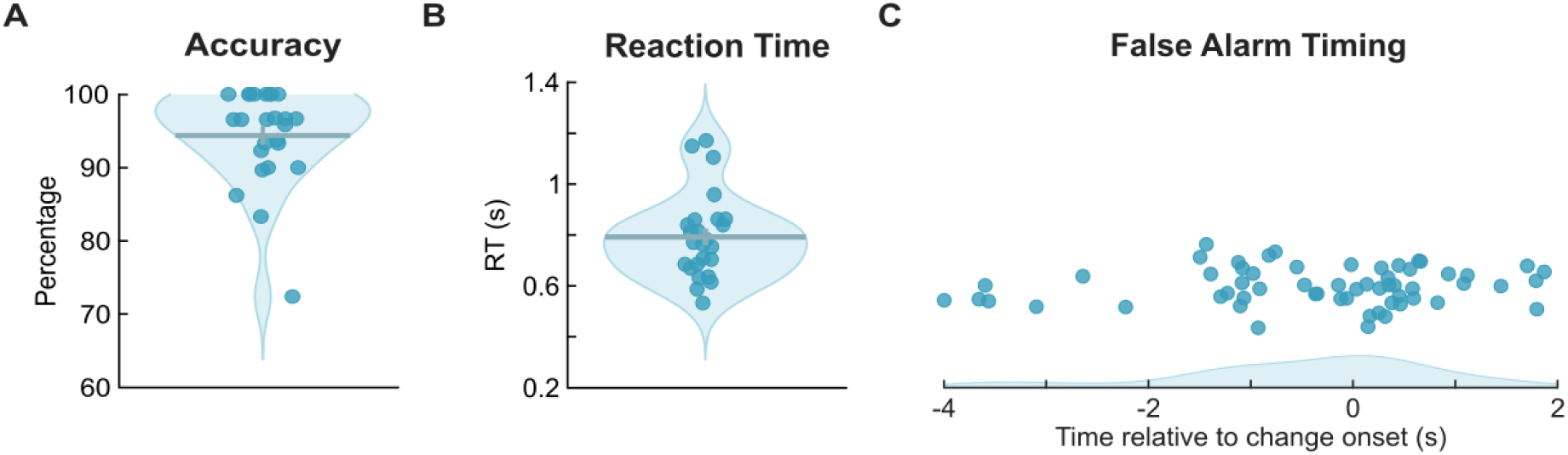
Decoy (Gap Detection) Task Performance. **[A]** Accuracy and **[B]** reaction time for the decoy task are shown across participants. **[C]** Within-trial distribution of false alarms (collapsed across all participants); each false alarm event is indicated by a dot. Change onset is at 0 s. False alarms were rare overall; when they did occur, they were largely confined to the interval from ∼2 s after scene onset (−2 s in the plot) to ∼1 s before scene offset. Shaded areas represent probability density. Dots represent individual participant data. Horizontal and vertical lines show mean and SEM, respectively.

Second, we determined whether the appearing source (change), was unintentionally eliciting these false alarms. Figure 2C plots the time course of false alarms collapsed across participants. These responses tended to occur partway through the scene, within the −2 to +1 s window relative to change onset. In the post-change period, false alarms were slightly more frequent during NC scenes: 58.62% of events (17/29) occurred in NC trials. Overall, we found that participants successfully performed the decoy task, engaging with the auditory stimuli, and showed no systematic evidence that gaps were confused with source appearance.

### 3.2. Occulomotor metrics reveal autonomic responses to task-irrelevant changes in complex scenes

To examine general response dynamics to source appearance, we first collapsed across all location conditions. Figure 3A, B, C shows group-averaged microsaccade (MS), pupil dilation rate (PDR), and pupil diameter (PD) measures, comparing change vs. no change trials. The earliest effect was observed in the MS, which revealed robust MSI. A significant divergence from the NC condition emerged at approximately 85 ms, with MSI peaking around 300 ms post-change.

**Figure 3.**
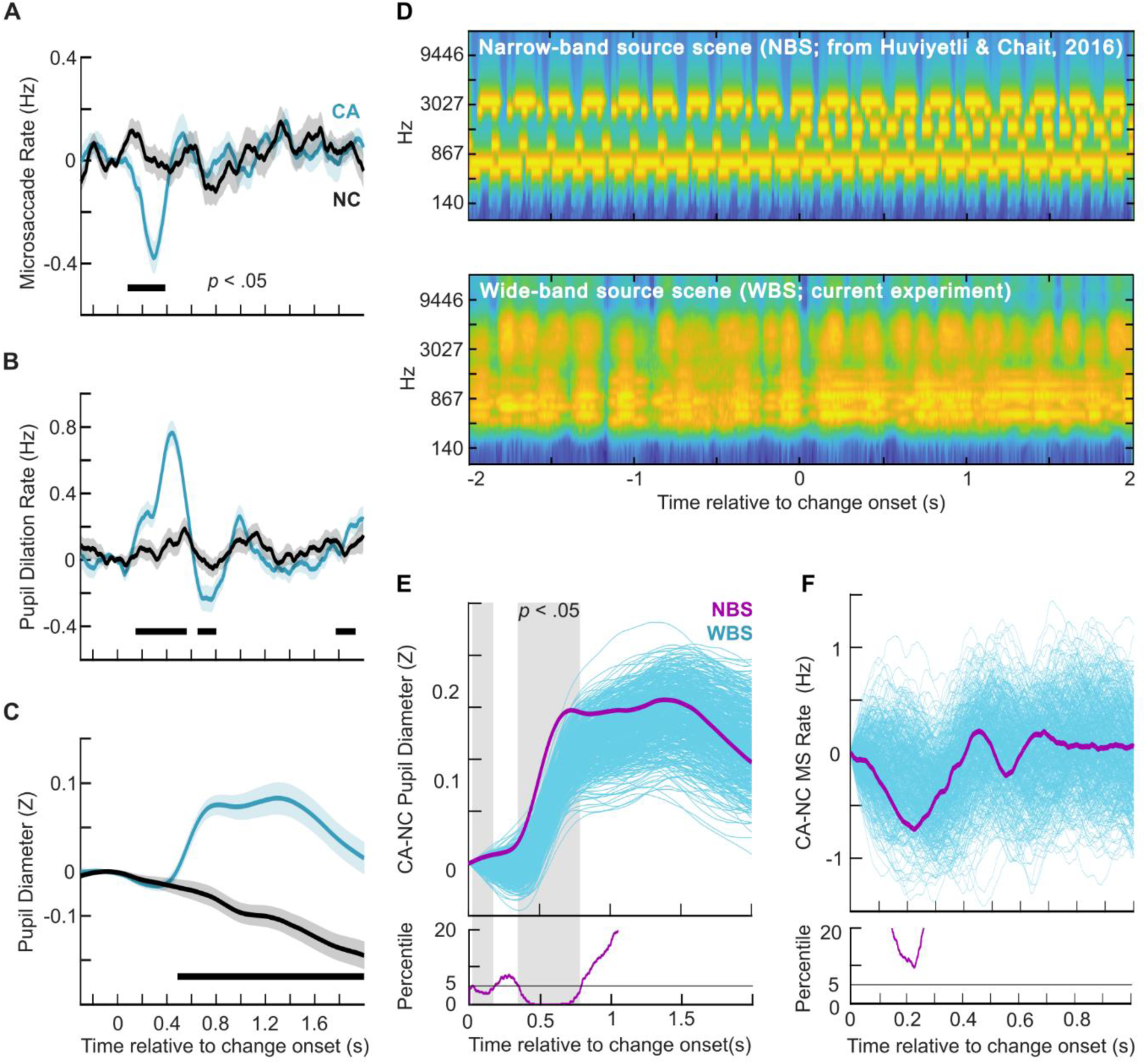
Change evoked ocular responses collapsed across conditions. **[A]** Microsaccade (MS) rate. **[B]** Pupil dilation rate (PDR). **[C]** Pupil dilation (PD) response. Shaded areas indicate SEM. Vertical bars (black) indicate significant differences between the conditions (p < .05; see Statistical Analysis). [D-F] **Comparison of PD and MS responses in spatialized wide-band scenes (present study) versus non-spatialized narrow-band scenes (Huviyetli and Chait, 2026). [D]** Example spectra for narrow-band source scenes (NBS; top) and wide-band source scenes (WBS; bottom). In these examples, a change (manifested as the appearance of a new source) is introduced partway through the scene (0 ms). Audio examples are at DOI:10.5522/04/32113429 **[E]** Change-evoked PD responses. The purple trace shows NBS data (N = 30; ∼30 trials per participant). Blue traces show resampled WBS data from the present experiment: on each iteration, 30 trials were randomly selected (from a total of 150 change trials) per participant and averaged across participants. The lower panel plots, at each time point, the percentage of iterations in which the NBS trace exceeds the WBS traces; the shaded region marks intervals in which fewer than 5% of WBS iterations exceed the NBS trace. This demonstrates that PD to changes in WBS is delayed by ∼150ms relative to simpler NBS **[F]** Same as [E], but for MS responses. No significant difference was found between NBS and WBS

PDR (see Methods), an index linked to phasic locus coeruleus activity (Joshi et al., 2016), began to increase from approximately 200 ms after the change and peaked at around 400 ms. This was followed by a significant phasic arousal response in PD, beginning approximately 400 ms after change onset. The temporal profile of this response is consistent with the double-peaked dynamics previously reported for tone-pip scenes (Huviyetli & Chait, 2026), which were interpreted as reflecting an initial arousal response to the change-related transient, followed by a second processing stage associated with pop-out of the appearing source (see also Sohoglu & Chait, 2016). Together, these findings indicate a progression of rapid reorienting responses to task-irrelevant changes in the auditory scene.

Next, we assessed how these responses to ecologically inspired stimuli relate to previously characterized ocular responses to simpler scene changes. Specifically, we compared the present PD and MSI responses with those reported by Huviyetli and Chait (2026), who used a similar paradigm but with scenes composed of isochronous narrowband sources (pure-tone streams). Example spectra for the two scene types are shown in Figure 3D. In both cases the scene contains a change - an abrupt appearance of a new source partway through the scene. This change is visually salient in the narrowband (tone-stream) spectrogram, but is much less apparent in the wideband scene (bottom), where energy is distributed across frequency and overlaps more strongly with the ongoing background. Importantly, despite the differences in spectral overlap, the changes are readily audible in both stimulus classes (audio examples: 10.5522/04/32113429). Relative to the narrow-band source scenes, the PD response observed here was delayed by approximately 150 ms, suggesting slower PD-linked arousal dynamics for changes embedded in more complex acoustic scenes. By contrast, no clear difference was observed in the MS response. As shown in Figure 3F, however, this may reflect the relatively high variability in MS responses in the present experiment. The mean latency of the MSI trough was 350 ms, compared with 200 ms for the simpler scenes reported by Huviyetli and Chait (2026). This increased variability is likely attributable to the absence of clear change-related frequency transients in the complex scenes used here.

### 3.3. Microsaccade dynamics are broadly similar across change locations

Figure 4 shows MS responses by spatial location. Changes in all locations elicited a pronounced MSI, with significant difference from the NC condition emerging from ∼200 ms post onset (Figure 4A) in all conditions and with a trough around 300–400 ms after change onset.

**Figure 4.**
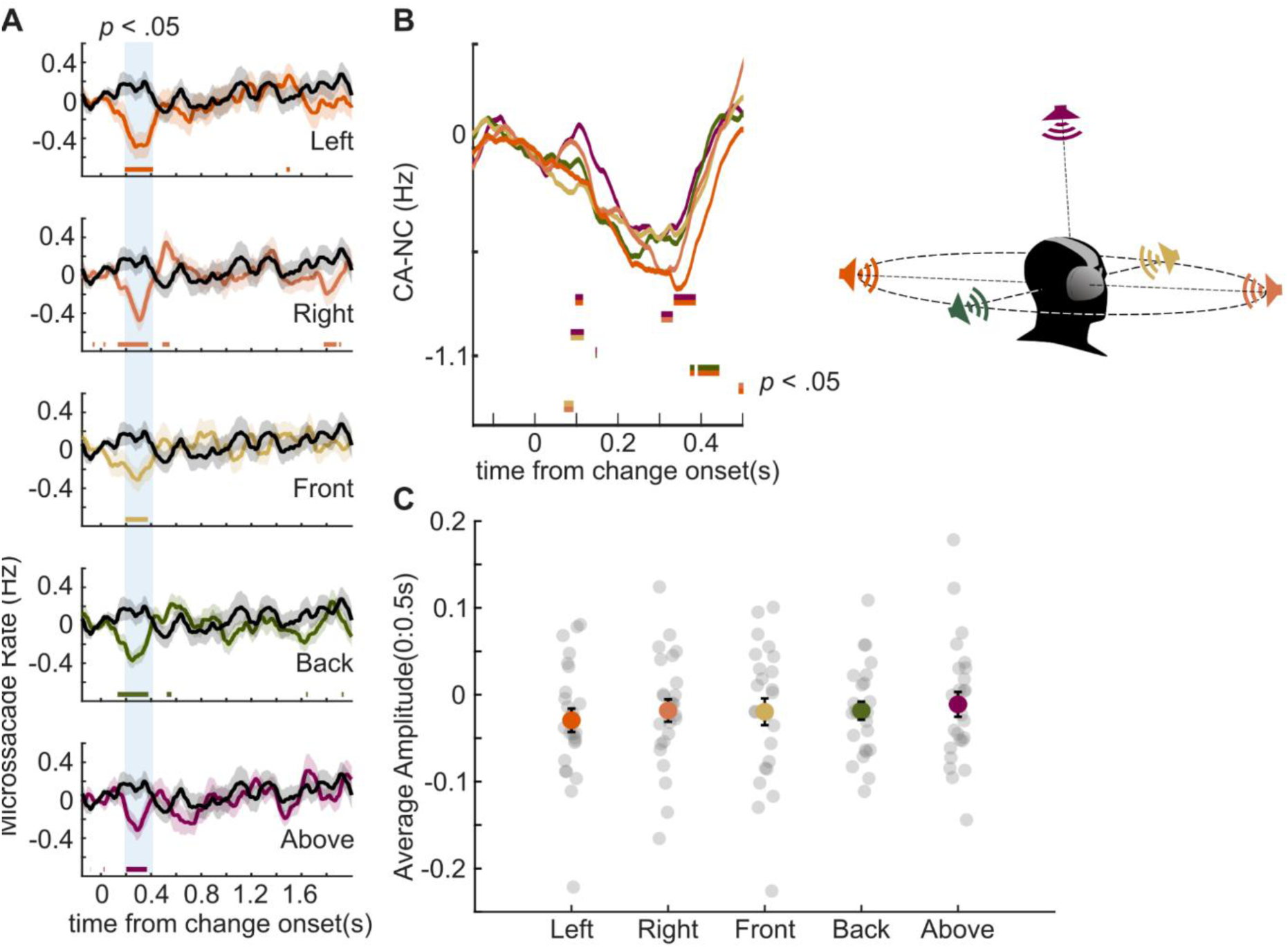
Microsaccade rate as a function of change location. **[A]** Baseline-corrected, change-evoked MS rate time courses for each appearing source location, expressed relative to the no-change control (NC; black). Coloured horizontal bars denote time intervals with significant differences between conditions; shaded regions around traces indicate ±SEM. Blue shading highlights the significant difference in the Left condition and is included to help readers compare significant effects across locations, indicated by the coloured horizontal lines. **[B]** Difference waves (CA − NC) for each location focusing on the initial MSI response. Horizontal bars mark significant differences; bar colors indicate the pairwise comparison being tested. **[C]** Mean response amplitude (0–500 ms) for the difference waves in [B]. Gray points show individual participants.

When comparing the MS responses for each condition, no differences between appearing source locations were seen during the initial phase (downward slope) of the MSI response. This suggests that MS-linked bottom-up attentional capture was similar across conditions. Brief differences (e.g., Left vs. Back/Above) are observed during the recovery phase (upwards slope of the MS time series) after about 400 ms post change onset. In addition to the time resolved analyses, we also quantified the differences between conditions by computing the mean CA–NC difference over the 500 ms post-change window (Figure 4C). A repeated-measures ANOVA showed no reliable effect of location, *F*(4, 96) = 0.74, *p* = 0.60.

### 3.4. Pupil dilation differs by appearing source location

To characterise the initial dynamics of the phasic pupil response, we analysed the PDR. PDR is an instantaneous index of phasic arousal that quantifies pupil responsivity and the incidence of pupil dilation events, irrespective of pupil size, and it has been linked to firing rates in the locus coeruleus (LC) and inferior colliculus (IC) (Joshi et al., 2016). Figure 5A shows PDR results stratified by the location of the appearing source. We observed a clear divergence from the NC condition at every location. Significant effects emerged from approximately 350 ms post-change onset for the Left, Right, Front, and Back conditions, whereas the Above condition showed a later significant divergence from NC, beginning at around 400 ms.

**Figure 5.**
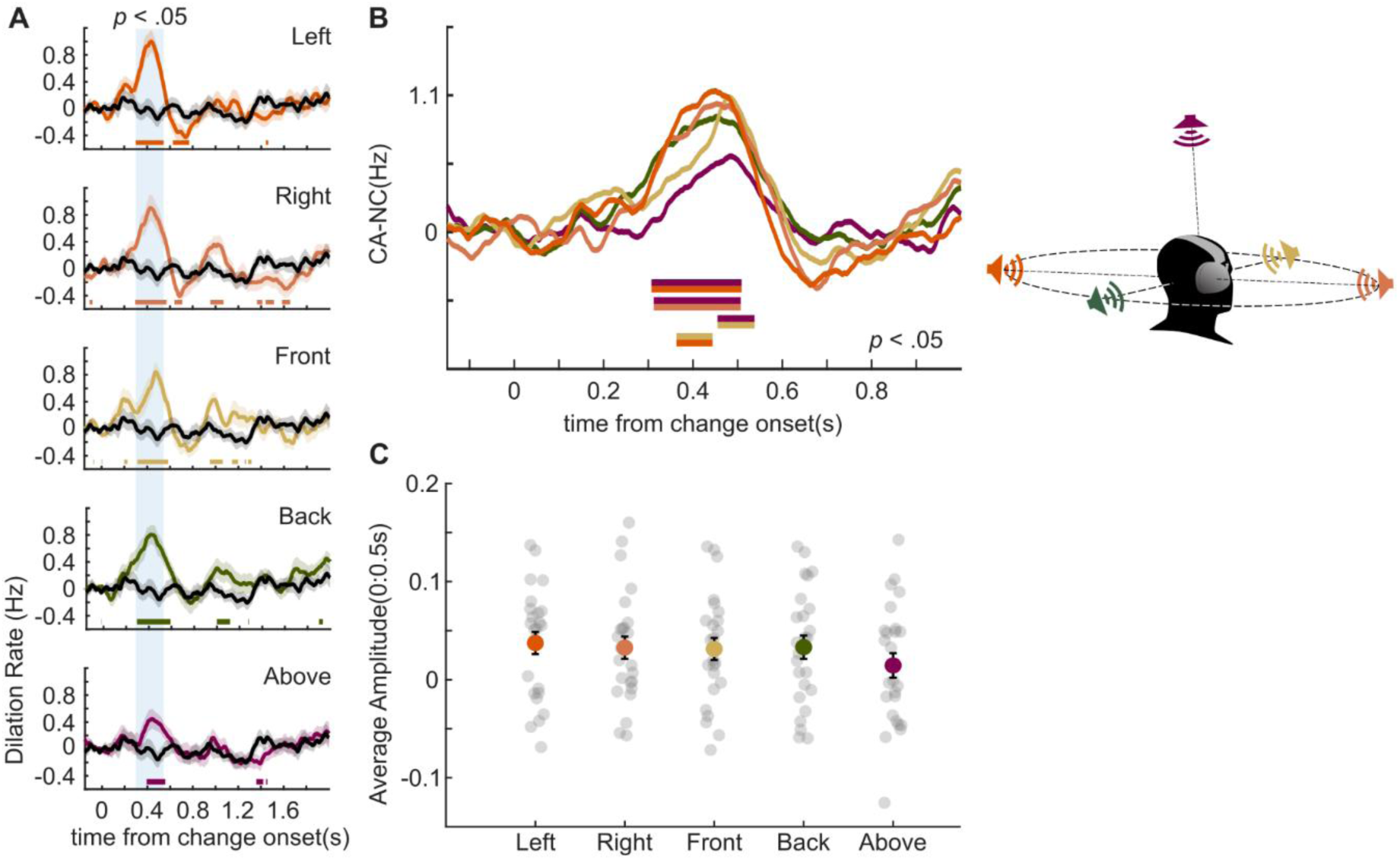
Pupil dilation rate as a function of change location. **[A]** Baseline-corrected, change-evoked pupil-dilation rate (PDR) time courses for each appearing source location, expressed relative to the no-change control (NC; black). Coloured horizontal bars denote time intervals with significant differences between conditions; shaded regions around the traces indicate ±SEM. Blue shading highlights the significant difference in the Left condition and is included to help readers compare significant effects across locations, indicated by the coloured horizontal lines. **[B]** Difference waves (CA − NC) for each location. Horizontal bars mark significant differences; bar colors indicate the pairwise comparison being tested. **[C]** Mean response amplitude (0–500 ms) for the difference waves in [B]. Gray points show individual participants.

Direct comparison of the condition traces (Figure 5B), plotted as the difference between each condition and NC, further suggest that the Front and Above conditions showed a delayed rise in PDR relative to the other conditions. Although the Front condition eventually reached a peak similar to those of the Left, Right, and Back conditions, the Above condition remained lower than the others throughout. However, when we assessed window-averaged PDR amplitude over the first 500 ms post-change, a repeated-measures ANOVA revealed no significant effect of location (F(4, 96) = 0.63, p = .550).

We then examined the PD response (Figure 6A-D). All conditions diverged from NC at approximately the same time, around 400 ms post-change onset (Figure 6A), suggesting that change events engaged arousal-related responses with a similar onset latency across locations. The conditions differed, however, in response amplitude, indicating location-dependent variation in evoked arousal. In particular, the differences between the Above condition and the other locations that were apparent around the peak of the PDR response were also evident in the subsequent dilation dynamics.

**Figure 6.**
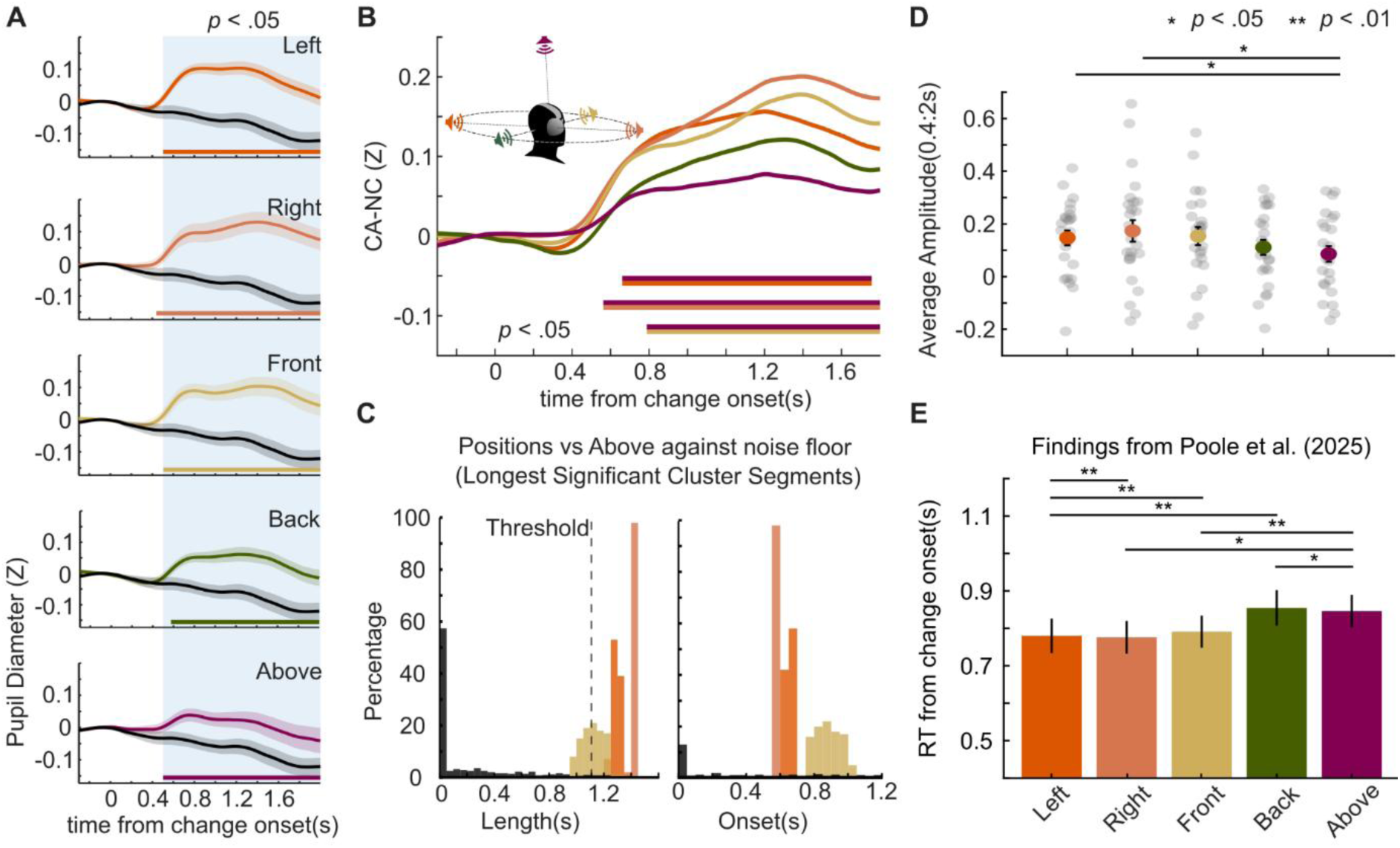
Pupil dilation responses as a function of change location. **[A]** Baseline-corrected, change-evoked pupil-dilation time courses for each location condition, expressed relative to the no-change control (NC; black). Coloured horizontal bars denote time intervals with significant differences between conditions; shaded regions around traces indicate ±SEM. Blue shading highlights the significant difference in the Left condition and is included to help readers compare significant effects across location conditions, indicated by the coloured horizontal lines. **[B]** Difference traces (CA − NC) for each location condition. Horizontal bars mark significant differences; bar colors indicate the pairwise comparison being tested. **[C]** Resampling (bootstrap) analysis. Black distributions define the noise floor, estimated by repeatedly resampling two sets of 30 trials from the NC condition and comparing them using the same procedure as for the main contrasts (see Methods). For each iteration, we retained either the longest significant interval (left) or the onset latency of longest significance (right). The dashed line indicates the 5th percentile of the noise-floor distribution. Colored distributions show corresponding metrics for contrasts of Left, Right, Front, and Back against Above. Left/Right/Front differ reliably from Above, whereas Back does not. **[D]** Mean response amplitude (0–2 s) for the difference waves in [B]. Gray points show individual participants. Significant differences are observed between Left and Above and between Right and Above. **[E]** Group averaged reaction time to detect an appearing source among five background maskers from Poole et al. (2025a), where listeners actively listened to similar stimuli presented via speaker array. Back and Above were associated with slower reaction time (RT) compared to the other location conditions.

Direct comparisons of the difference traces (CA–NC; Figure 6B) indicated location-dependent modulation of phasic arousal: PD responses were significantly larger for changes presented in the Front, Left, and Right locations than for changes presented Above. This difference emerged from approximately 600 ms post-change onset and persisted until stimulus offset. Figure 6C shows direct comparisons between the Above condition and the Front, Left, Right, and Back conditions, evaluated against a noise floor estimated by resampling the NC condition. This analysis indicates no reliable difference between Back and Above, with the observed effect indistinguishable from the noise floor. In contrast, Above differed significantly from Front, Left, and Right. The observed distinction between Front/Left/Right and Above/Back is consistent with previously reported delays in reaction times for detecting sounds appearing behind and above the listener (Poole et al., 2025, reproduced in Figure 6E).

To quantify these effects more directly, we computed the mean CA–NC difference within a 400–2000 ms post-change window (Figure 6B, bottom). A repeated-measures ANOVA revealed a significant main effect of location, F(4, 96) = 3.09, p = .031. Post-hoc pairwise comparisons confirmed significant differences between Above and Left, t(24) = 2.56, p = .017, and between Above and Right, t(24) = 2.61, p = .015, with a marginal difference between Above and Front, t(24) = 2.56, p = .057. Overall, these results indicate that phasic PD varies as a function of change location, with broadly similar responses for Left, Right, and Front, but a reduced response for Above.

Overall, the three ocular response analyses revealed a gradual divergence across conditions. No differences were apparent during the initial MSI response. However, during the rebound phase at approximately 300 ms, a difference emerged between the Left/Right conditions and the Front/Back/Above conditions, with the latter three exhibiting a shallower MSI trough. Around 400 ms post-onset, during the PDR peak period corresponding to the initial phasic arousal response, the Above condition showed a reduced response, which persisted into the PD period.

### 3.5. Behavioral localization reveals uncertainty for sources appearing from the front, back, or above

Figure 7 summarizes performance on the localization task (N = 25), administered at the end of the session. Participants listened to CA scenes (identical to those presented during the main experiment) and, for each change event, reported the perceived source location along with their confidence in that judgment. Panels A–C show results for the HRTF-based scenes.

**Figure 7.**
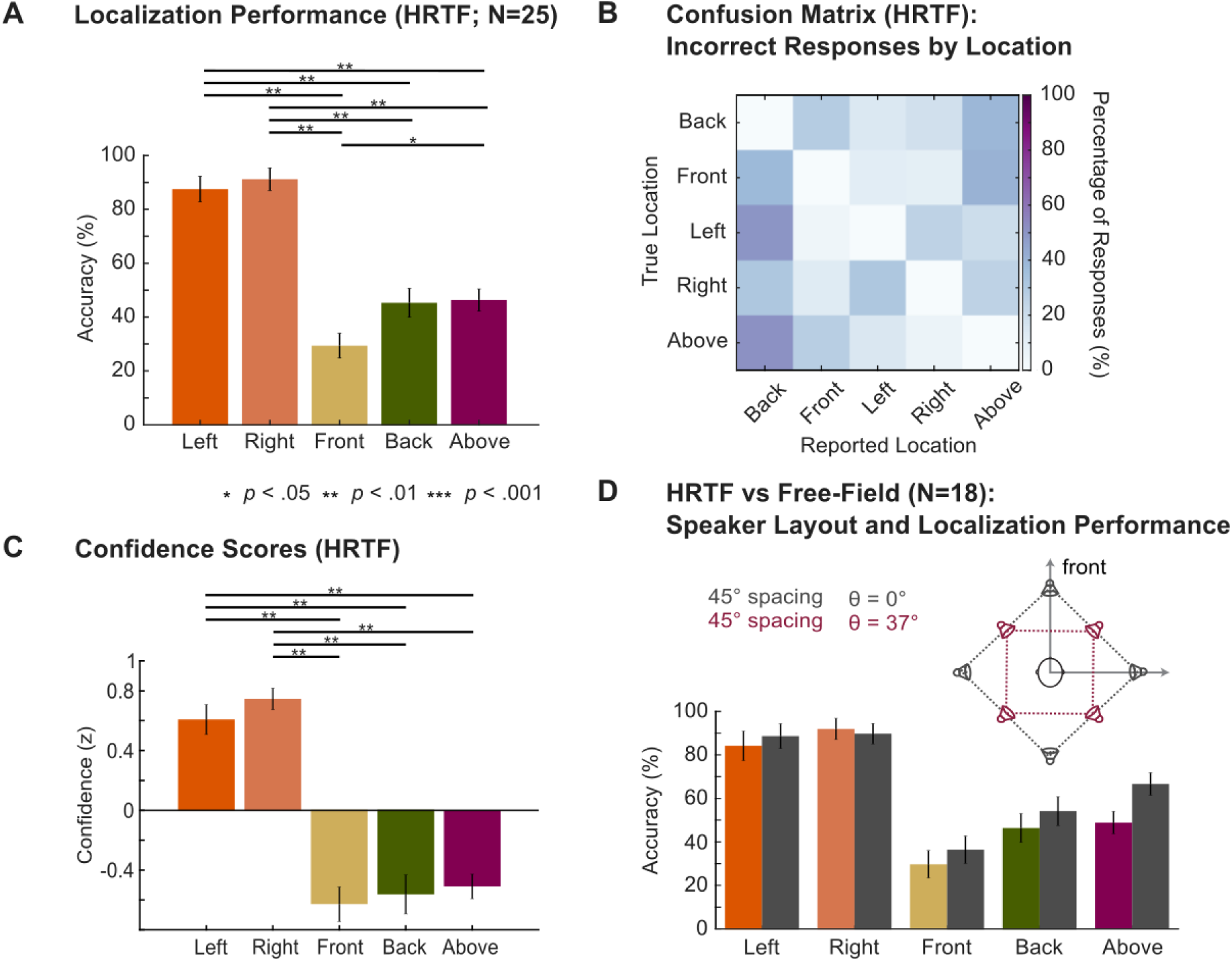
Behavioral localization task performance conducted following eye tracking. **[A]** Localization performance for HRTF (headphone-based) rendering of sound sources (N=25). Vertical error bars represent SEM. Horizontal lines mark significant pairwise comparisons . **[B]** Localization confusion matrix reporting confusion patterns for incorrect responses (hence values along the diagonal are 0).**[C]** Confidence scores. Individual z-scores were calculated against each participant’s average confidence score across conditions.**[D]** Comparison of localization performance between free-field (dark grey bars) and HRTF-rendered scenes (N=18).The free-field speaker layout is provided in the inset, with the speakers coloured by the elevation angle (black = 0°, purple +37°). The ear level appearing sources were presented as real loudspeaker sources. The amplitude panning method (Pulkki, 1997) was used to present the Above appearing source and positions of background sources identical to those in the HRTF version (see Figure 1).

Lateral positions were identified with high accuracy(Left=87.2%, Right=91.2%), whereas Front/Back/Above locations were localized much less accurately(29.6%, 45.6%, 49.6%, respectively), with frequent confusions among these categories. A Friedman test showed a significant effect of location on accuracy scores, χ²(4) = 65.28, p < .001. Post-hoc comparisons (Wilcoxon signed-rank test with Bonferroni correction) revealed significant differences for Left compared to Back, *Z* = 4.17, *p* < .001, Front, *Z* = 4.18, *p* <.001, and Above, *Z* = 3.98, *p* < .001 as well as Right compared to Back *Z* = 4.32, *p* < .001, Front, *Z* = 4.33, *p* < .001, and Above, *Z* = 4.17, *p* < .001. We also observed significant differences between Above and Front, *Z* = 2.90, *p* = .037. This pattern is mirrored in participants’ self-reported confidence, which was notably lower for Front/Back/Above conditions. Identical analysis showed significant effect of location, χ²(4) = 65.28, p < .001, with post-hoc tests comparison uncovering significant differences for Left compared to Back, *Z* = 3.62, *p* = .003, Front, *Z* = 3.98, *p* <.001, and Above, *Z* = 3.75, *p* = .002 as well as Right compared to Back *Z* = 3.94, *p* < .001, Front, *Z* = 4.04, *p* < .001, and Above, Z = 3.91, *p* < .001.

To investigate whether these effects reflect general auditory localization and are not specific to the HRTF rendering, in a subset of participants (n = 18; Figure 7A), we compared localization performance for HRTF (headphone-based) rendering versus free-field presentation (see Methods; Figure 7D). The overall pattern was similar across renderings, with reduced accuracy for Front/Back/Above locations. A repeated-measures ANOVA showed a significant main effect of location, *F* (4,68) = 47.6 , *p* < .0001, but no main effect of rendering (HRTF vs Free-Field), *F* (1,17) = 2.69, *p* = .12 and no interaction, *F* (4,68) = 0.84 , *p* = 0.504. Overall these results demonstrate evidence for midline confusion in both renderings, with Front being somewhat worse than Back/Above. It is noteworthy that the pattern of perceptual confusion exhibited by listeners differs from the pattern of PD-linked arousal or MS-linked attentional capture (see further discussion below).

## 4. Discussion

This study demonstrates that abrupt, task-irrelevant changes in complex spatialized auditory scenes elicit rapid, stereotyped reorienting responses in the oculomotor–autonomic system, even when those changes are not behaviorally relevant. Across all change locations, we observed robust microsaccadic (MS) inhibition and phasic pupil dilation (PD), consistent with automatic detection of auditory novelty. Previous work has shown that both MS and PD responses are modulated by the bottom-up perceptual salience of sound events (Bianco et al., 2021; Huviyetli & Chait, 2026; Zhao et al., 2019). We therefore expected that, if spatial factors influenced bottom-up perceptual salience at the level indexed by MS or PD, this would be reflected in our physiological measures.

Consistent with earlier reports of dissociations between MS and PD responses (Huviyetli & Chait, 2026; Liu & Chait, 2026; Zhao et al., 2024), these two markers differed in their sensitivity to spatial factors in the present data. The initial stage of MSI was largely invariant across change locations whereas pupil-linked arousal varied as a function of the location of the appearing source. Notably, however, neither physiological measure mirrored the pattern of behavioral confusion during active localization. In the behavioral data, participants showed high confidence for Left and Right locations, but comparably reduced confidence and accuracy for Front, Back, and Above sources. By contrast, during ocular recording, when the scene changes were task-irrelevant, changes originating from the Front elicited responses similar to those evoked by Left and Right sources, whereas changes from Above (and to some extent those located in the Back) were associated with reduced phasic arousal.

Taken together, these findings reveal a dissociation between the perceptual confusion expressed during active localization judgments and the autonomic responses observed during passive, non-target listening.

### 4.1. PD, PDR, and MSI reveal automatic re-orienting in complex scenes

The group averaged microsaccade data revealed a clear MSI response emerging rapidly (∼85 ms) and peaking at ∼300 ms post-change. Pupil data revealed an increase in PDR starting ∼200 ms and peaking around ∼400 ms which was accompanied by a significant phasic PD response beginning ∼450 ms after change onset. This temporal progression, in line with that seen in other reports ( Huviyetli & Chait, 2026; Liu & Chait, 2026; Zhao et al., 2024) reveals a multi-component orienting response to scene change: a fast oculomotor interruption (MSI) and a slightly later autonomic arousal component (PD/PDR).

Comparisons with a closely related paradigm using isochronous pure-tone scenes (Huviyetli & Chait, 2026) suggest that the PD response in the present complex scenes is delayed by ∼150 ms. A similar delay was observed when comparing EEG responses: change-evoked responses in wideband, spatialized scenes emerged ∼150 ms later (∼200 ms post-change; Poole et al., 2025a) than responses in scenes comprising narrowband streams (∼50 ms post-change; Sohoglu & Chait, 2016a). In contrast, MSI timing was comparable across paradigms. A parsimonious interpretation is that complex scenes impose additional sensory analysis or object-formation demands before a change event is fully evaluated as “novel/salient” at the level that drives EEG and pupil-linked arousal. Pupil dilation reflects a comparatively integrative process, potentially incorporating salience evaluation, uncertainty, and global state modulation, whose latency may therefore be sensitive to scene structure. MSI, by contrast, may index an earlier, more stereotyped interrupt signal triggered by deviance detection that is less dependent on the richness of scene content, consistent with the idea that microsaccadic suppression is a fast, modality-general reorienting mechanism.

### 4.2. Behaviour confirms spatial uncertainty outside the lateral axis

Localization performance measured after the eyetracking experiment revealed that participants could reliably identify lateral positions, but showed marked uncertainty for Front/Back/Above locations, characterized by frequent confusions and reduced confidence. This aligns with known ambiguities in spatial hearing (Riedel, Frank, & Zotter, 2025; Zhang & Hartmann, 2010), particularly for elevation and front–back discrimination under constrained cue conditions, and is consistent with the idea that the perceptual representation of these locations is less stable and more probabilistic than for lateral positions (Barumerli et al., 2022; Ege, Opstal, & Van Wanrooij, 2018; Poirier-Quinot, et al., 2022, Reijniers et al., 2025).

Critically, these behavioural judgment data provide important context for interpreting the physiological differences observed across locations. In principle, reduced certainty and/or lower salience for particular locations could plausibly attenuate arousal responses. However, the pattern of location-dependent modulation observed in the ocular data did not align with the behavioural localisation data. Although PD responses were lowest for the “Above” condition, and to some extent for “Back”, responses to “Front” did not differ from those to the lateral positions. Instead, the PD data were more consistent with behavioural change-detection findings reported by Poole et al. (2025a, 2026; see Figure 5E), in which participants indicated the presence of a change irrespective of its location. In those studies, performance was poorer for the Above and Back conditions than for Front/Left/Right.

What appears clear from the ocular data, as discussed further below, and from the previous EEG findings of Poole et al. (2025a, 2026), is that there is no evidence for condition differences in the early response period. These location-based behavioral biases therefore likely arise later in the processing stream, after approximately 200 ms following change onset. One possibility is that the localisation confusion between Front, Back, and Above, observed when participants are required to determine **where** a scene change has occurred, reflects a late-stage process that is independent of change detection itself, that is, of deciding **whether** a change has occurred. Alternatively, ambiguity among the Front, Above, and Back locations may operate in tandem with a separate bias favouring the Front (Best, Boyd, & Sen, 2023; Montagne & Zhou, 2018; Pomper & Chait, 2017), thereby producing the pattern observed in the PD data and in behavioural change-detection performance.

### 4.3. Microsaccades as a location-invariant marker of change detection

MSI, during its initial phase, showed little dependence on change location. All locations elicited a pronounced trough around 300–400 ms, and window-averaged metrics did not differ reliably across conditions. This pattern suggests that MSI reflects a relatively generic novelty or interruption response that is less sensitive to the specific spatial attributes of the deviant event.

These observations are consistent with findings from an EEG experiment using the same stimuli (Poole et al., 2025a), in which no differences were found between location conditions. In that study, EEG responses to scene changes emerged at around 200 ms, broadly matching the timing of the MSI effects observed here. The present findings are also consistent with Zhao et al. (2024), who examined the effects of top-down attention on MSI responses to simple acoustic events embedded in either an attended or an ignored stream. They reported that the initial phase of the MSI response was unaffected by attention, whereas attention modulated the duration and rebound of the response. This closely resembles the pattern observed here, with location differences emerging only at a later stage, and supports the view that the initial MSI response reflects a reflexive change detection process.

Although it is difficult at this stage to exclude the possibility that the absence of location effects in early MSI and EEG reflects signal noise, several arguments suggest that this is unlikely. First, the EEG effect was robust even when combining data from 62 participants - a large number with which we would typically expect sufficient power to detect between-condition differences. Second, previous work has shown that subtle differences in MSI dynamics are detectable with the current general paradigm (in terms of trial numbers and sample sizes; Huviyetli & Chait, 2026; Zhao et al., 2024).

### 4.4. Location-dependent modulation of pupil responses

A key novel finding is that PD responses, unlike MSI, varied with change location, but did not fully mirror the behavioral localization pattern. Specifically, the Front–Back–Above confusions commonly observed in behavioral localization responses were not evident in pupil responses recorded during passive listening. Pupil responses to Front were comparable to those for Left and Right, with attenuation primarily for Above (and, to a lesser extent, Back). In particular, Front, Left, and Right elicited larger phasic dilation than Above, with a significant main effect of location in the 400–2000 ms window and post-hoc differences driven mainly by the reduced response for Above. PDR showed a qualitatively similar separation around its peak. Importantly, the latency of the PD response (Figure 5) did not vary across spatial locations, with differences emerging only in response magnitude. This is consistent with an account, also supported by the MSI data above, that the initial detection of the change occurred at a similar time across conditions, and that the observed PD effects were driven by differences in the strength of arousal-related modulation rather than by differences in response latency.

Several non-mutually exclusive mechanisms could account for reduced pupil responses to Above/Back. First, changes from above may be perceptually less reliable, regarding the location of the appearing source, under the present HRTF conditions, leading to reduced confidence and/or weaker evidence accumulation about the event’s spatial attributes; pupil-linked arousal has been repeatedly associated with uncertainty and decision-related processes (Fan et al., 2023; Milne et al., 2021; Urai et al., 2017; Vincent et al., 2018; Zhao et al., 2019), and may therefore scale with the strength or clarity of the inferred change. Second, above sources may have been less salient due to inherent differences in the spectral information from this location (e.g., less distinctive spectral profiles differentiating above from other spatial positions), producing a weaker orienting response. Third, listeners may implicitly treat certain spatial regions (e.g., lateral/front) as more behaviourally relevant or ecologically probable, and pupil-linked arousal could reflect such implicitly learned priors about potential importance, therefore producing stronger arousal modulation for Left/Right/Front relative to Back/Above.

Distinguishing between these accounts will require future manipulations that independently vary acoustic salience, spatial certainty, and task relevance. Notably, however, both MSI responses (see above) and EEG responses (Poole et al., 2025a) show no differences between location conditions. Furthermore, a numerical HRTF analysis reported in a related study (Poole et al., 2026) found that acoustic shadowing was minimal between Front and Back locations and with small frequency-specific RMS differences of approximately 5 dB. This suggests that a simple explanation based on acoustics is unlikely.

Notably, the pattern of reduced pupil responses for Above (and Back) aligns with behavioural findings reported in both normal-hearing listeners (Poole et al., 2025a) and hearing-impaired listeners (Poole et al., 2026). In those experiments, based on the same stimuli as used in the present experiment, participants engaged in an active change detection task in free-field, where they were asked to indicate when they perceived that a change occurred in the scene but there was no localization response made. Change detection performance was consistently poorest for sources in Back and Above conditions (see also Figure 5E, above). Together this would indicate that reduced behavioral change detection (though not tested directly in the present experiment) for Above/Back is a consequence of a reduced engagement of the LC-NE arousal system in those conditions.

### 4.5. MSI and PD dynamics reveal the emergence of a perceptual behavioral bias

Taken together, the emerging picture is that early change-detection responses in the auditory cortex, indexed by EEG and associated MSI responses linked to attentional orienting, are largely insensitive to the spatial location of the change. Instead, a bias against Above and Back locations appears to arise later in the processing stream, influencing arousal responses (in naïve, distracted listeners), and behavioural change detection performance (as reported in Poole et al., 2025a; 2026), and potentially related to perceptual localization confusion (as seen here; Figure x).

The finding that PD and MS are differentially modulated is consistent with a growing body of evidence suggesting that these measures provide complementary indices of processing demands (Contadini-Wright et al., 2023; Huviyetli & Chait, 2026; Liu & Chait, 2026; Zhao et al., 2024). PD has been linked more closely to arousal-related modulation, whereas MS is increasingly interpreted as an indirect index of attentional resource allocation. Although arousal and attention both form part of the orienting response and are supported by partially overlapping brainstem circuitry, including regions such as the LC and IC, they appear to be at least partly dissociable with the sequence potentially reflecting the initial stages of bottom up attentional capture by a new event (MSI) which then triggers an arousal response (PDR and PD).

The present findings extend this emerging view by showing that location-related perceptual biases are not apparent at earlier stages of the processing hierarchy, as indexed by EEG and microsaccadic responses, but are instead expressed in pupil dilation. More broadly, these results suggest that listening challenges in complex spatialized scenes can be investigated by tracking how these response components vary across task-relevant manipulations, and how they are altered in hearing-impaired populations, in whom attentional capture and/or arousal regulation may be disrupted. By jointly leveraging MS and PD as complementary markers of attentional orienting and arousal, this approach offers a scalable framework for examining how hearing impairment affects the dynamic updating of auditory scene representations and, consequently, the maintenance of situational awareness in real-world listening.

## Acknowledgements

The work was funded by the Demant foundation (project ANTHEA). The funders had no role in study design, data collection, and analysis, the decision to publish, or preparation of the manuscript.

## Data sharing

The data reported in this manuscript alongside related information will be available at DOI: 10.5522/04/32122729

